# The tetrameric assembly of 2-aminomuconic acid dehydrogenase is a functional requirement of cofactor NAD^+^ binding

**DOI:** 10.1101/2021.07.15.452591

**Authors:** Qiuli Shi, Yanjuan Chen, Xinxin Li, Hui Dong, Cheng Chen, Zhihui Zhong, Cheng Yang, Guangfeng Liu, Dan Su

## Abstract

The bacterium *Pseudomonas* sp. *AP–3* is able to use the environmental pollutant 2-aminophenol as its sole source of carbon, nitrogen, and energy. Eight genes (amnA, B, C, D, E, F, G, and H) encoding 2-aminophenol metabolizing enzymes are clustered into a single operon. 2-aminomuconic 6-semialdehyde dehydrogenase (AmnC), a member of the aldehyde dehydrogenase (ALDH) superfamily, is responsible for oxidizing 2-aminomuconic 6-semialdehyde to 2-aminomuconate. In contrast to many other members of the ALDH superfamily, the structural basis of the catalytic activity of AmnC remains elusive. Here, we present the crystal structure of AmnC, which displays a homotetrameric quaternary assembly that is directly involved in its enzymatic activity. The tetrameric state of AmnC in solution was also presented using small-angle X-ray scattering. The tetramerization of AmnC is mediated by the assembly of a protruding hydrophobic beta-strand motif and residues V121 and S123 located in the NAD^+^-binding domain of each subunit. Dimeric mutants of AmnC dramatically lose NAD^+^ binding affinity and enzyme activity, indicating that tetrameric assembly of AmnC is required for oxidizing the unstable metabolic intermediate 2-aminomuconic 6-semialdehyde to 2-aminomuconic acid in the 2-aminophenol metabolism pathway.

## Introduction

*Pseudomonas* is a common genus of bacteria that can grow on a variety of carbon, nitrogen, and sulfur sources and produce multiple secondary metabolites(1). This ability to degrade and produce a wide spectrum of compounds has led to a biotechnological interest in *Pseudomonas* sp. for bioremediation, as well as the production of polymers and low-molecular-weight compounds(2). Some *Pseudomonas* strains have been used to remove highly polluting compounds containing hard-to-degrade substances(3, 4). *Pseudomonas* sp. *AP–3* can metabolize 2-aminophenol and its ring-cleavage product, 2-aminomuconic 6-semi-aldehyde, which is further degraded via the meta-cleavage pathway(5). The pathway for 2-aminophenol degradation was proposed following the identification of metabolites and enzymatic assays of ring cleavage enzymes in *Pseudomonas* sp. *AP–3*(6). Eight enzymes translated from the AmnABCDEFGH gene cluster in the *Pseudomonas AP-3* strain have been identified to play a role 2-aminophenol metabolism. 2-aminophenol is converted to 2-aminomuconic 6-semialdehyde by the catalysis of 2-aminophenol 1,6-dioxygenase (AmnBA), which is then dehydrogenated to 2-aminomuconic acid in the presence of nicotinamide adenine dinucleotide (NAD+) by 2-aminomuconic 6-semialdehyde dehydrogenase (AmnC). Then, 2-aminomuconate deaminase (AmnE) deaminates the metabolite to form 4-oxalocrotonic acid, which is decarboxylated to 2-oxopent-4-enoic acid by 4-oxalocrotonate decarboxylase (AmnD). 2-oxopent-4-enoate hydratase (AmnF) hydrates 2-oxopent-4-enoic acid to 4-hydroxy-2-oxovaleric acid, which is finally converted into pyruvic acid and acetaldehyde by 4-hydroxy-2-oxovalerate aldolase (AmnG) and acetaldehyde dehydrogenase (AmnH)(7). The potential applications of the 2-aminophenol metabolism pathway warrant a detailed study and characterization of its component enzymes and their structures. In particular, one enzyme, AmnC, is responsible for oxidizing the unstable metabolic intermediate 2-aminomuconic 6-semialdehyde to 2-aminomuconic acid in the 2-aminophenol metabolism pathway. Based on sequence alignment, AmnC is a member of the aldehyde dehydrogenase (ALDH) superfamily, which is one of the most ancient protein superfamilies and is widely distributed in all three taxonomic domains (Archaea, Eubacteria, and Eukarya). The ALDH superfamily is responsible for oxidizing aldehydes to their corresponding carboxylic acids in a broad variety of metabolic pathways(8). ALDHs from *Pseudomonas* can be classified into 42 protein families, with a predominance of 14 families that contain 76.6% of all known ALDHs (9). Mechanistically, ALDHs use covalent catalysis to form a thiohemiacetal between a catalytic cysteine residue and the aldehyde substrate(10, 11). The thiohemiacetal is then oxidized to a thioester by hydride transfer to the coenzyme. Depending on the nucleophile used to break the thioester bond to release the acid product of the reaction, there are three types of ALDHs: hydrolytic, CoA-acylating, and phosphorylating. Specifically, hydrolytic ALDHs play important roles in catabolic and anabolic processes, as well as in the elimination of toxic aldehydes(12). CoA-acylating ALDHs allow for detoxification of aldehyde intermediates produced in degrading pathways and the formation of ATP through substrate-level phosphorylation using the newly formed acyl-CoA(13). Phosphorylating ALDHs are involved in proline synthesis (14). According to the classification of catalytic reactions, AmnC belongs to a group of hydrolytic ALDHs. In addition to metabolizing aldehydes, some ALDH isozymes have reductase and esterase activities and act as binding proteins for endogenous substrates. From a structural view, ALDHs share a highly conserved architecture to perform the oxidation of a broad range of aldehydes into corresponding carboxylic acids with the reduction of their cofactor, NAD^+^ or NADP^+^. The typical ALDH structure exhibits a three-domain architecture consisting of a Rossman dinucleotide-binding domain, an α/β-catalytic domain, and an oligomerization domain(15).

Currently, there are 49 structures available from members of the ALDH superfamily. Most adopt a tetrameric assembly; however, a small number of the resolved structures exhibit dimeric or hexameric assemblies. Despite this, the molecular mechanism by which ALDHs exert their catalytic activity through an oligomeric state remains elusive. Here, we determined the tetrameric structure of AmnC in *Pseudomonas sp. AP–3* using a combination of X-ray crystallography and small-angle X-ray scattering (SAXS). AmnC shares a highly conserved tetrameric conformation with the ALDH superfamily. Site-directed mutagenesis was used to identify the tetrameric contact interfaces, with a focus on the oligomerization domain and several highly conserved residues (S123G, H124G, and L425A) located in the catalytic domain of AmnC. Truncated versions of AmnC with a deletion of the oligomerization domain at residues Δ124-138 and Δ477-491 completely prevented tetramer formation in solution and also resulted in a loss of catalytic activity and NAD^+^ binding affinity, indicating that the tetramer interfaces significantly influence enzyme activity. In summary, we report the tetrameric assembly of 2-aminomuconic 6-semialdehyde dehydrogenase from *Pseudomonas sp. AP–3* and propose that homotetramerization is required for the enzymatic activity of AmnC through interaction with the cofactor NAD^+^.

## Results

### Catalytic activity of wild-type AmnC

The putative native substrate of AmnC, 2-aminomuconate-6-semialdehyde (2-AMS), is a proposed metabolic intermediate in the 2-aminophenol degradation pathway of *Pseudomonas* sp. *AP–3*. In the presence of NAD^+^ and AmnC, 2-AMS is oxidized to 2-aminomuconate (2-AM). However, 2-AMS can also spontaneously decay to picolinic acid and water with a half-life of 35 seconds in a neutral pH environment. Due to its instability, 2-AMS has not yet been isolated. 2-hydroxymuconate-6-semialdehyde (2-HMS) is a previously identified, stable alternative substrate in which a hydroxyl group replaces the amino group in 2-AMS to prevent cyclization in the presence of NAD^+^ and AmnC; 2-HMS is oxidized to α-hydroxymuconic acid (2-HM)(16). The activity of AmnC and mutants was routinely assayed by measuring the decrease in absorbance of 2-HMS at 375 nm in the presence of NAD^+^.

To obtain the stable alternative substrate 2-HMS, we used Fe^2+^ reconstituted 3-hydroxyanthranilate 3,4-dioxygenase (nbaC) from *Pseudomonas fluorescens* to catalyze the insertion of molecular oxygen to 3-hydroxyanthranilic acid (3-HAA); the resulting α-amino β-carboxymuconate ε-semialdehyde (ACMS) has an absorption maximum at 360 nm (Fig.1*A, B*). Lowering the pH below 3 led the compound to have an absorption maximum at 315 nm, which is the maximum absorbance of 2-HMS at pH 3. Before testing the catalytic activity of AmnC, we adjusted the pH of 2-HMS to 8.0, at which point its absorption maximum changes to 375 nm (Fig.1*A, C*) (16) (17). When 2-HMS was incubated with NAD^+^ and wild-type AmnC, the absorption maximum at 375 nm decreased rapidly and an absorption maximum at 335–340 nm appeared instead (Fig.1*D, E*), suggesting that the absorbance at 335–340 nm was due to the formation of NADH(18).

**Figure 1.**
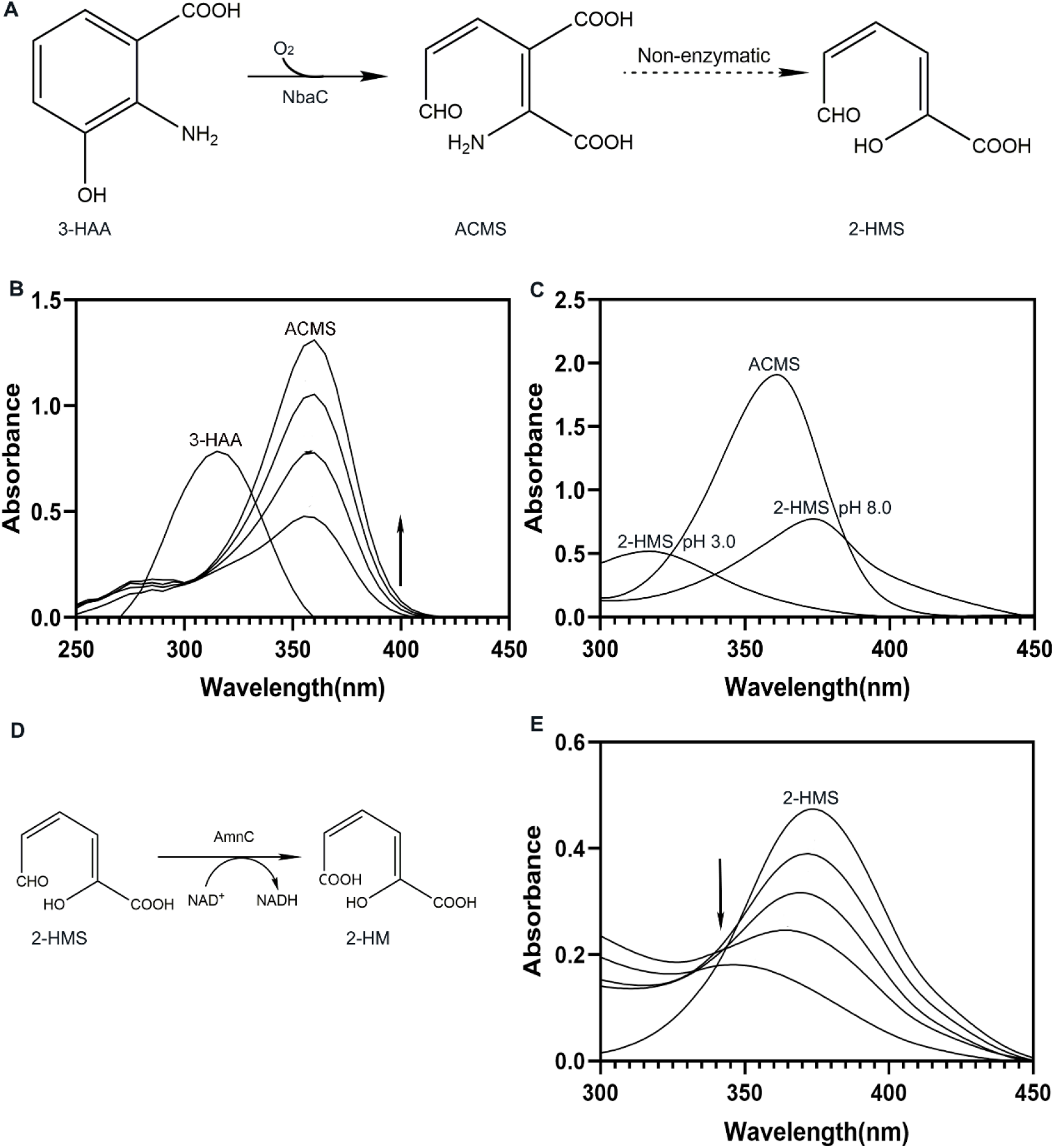
Catalytic activity of AmnC. (**A**) Reaction scheme showing the enzymatic generation of 2-HMS through a reaction catalyzed by NbaC. (**B**) Representative assay showing the NbaC-catalyzed conversion of 3-HAA (λmax 315 nm) to ACMS (λmax 360 nm). (**C**) Non-enzymatic reaction conversion of ACMS to 2-HMS (λmax 315 nm, pH3.0; λmax 375nm, pH8.0). (**D**) Reaction scheme showing 2-HMS oxidation by AmnC. (**E**) Representative assay showing the activity of AmnC on 2-HMS (λmax 375 nm).

### Structural overview of AmnC

To address the structural basis for the catalytic and regulatory mechanisms of AmnC in the meta-cleavage pathway, we determined the 1.9 Å resolution crystal structure of AmnC (residues 1-491), which was crystallized in the hexagonal space group P6_1_22 and contained two AmnC molecules in an asymmetric unit(19). The final refined model of AmnC contained residues 1-488 in both molecules and had an R_factor_ of 17.2% and an R_free_ of 20.7% (Table 1). Ramachandran plots showed that 99.5% of the residues were in the allowed region. To obtain additional information about the interfaces and likely biological assemblies of AmnC multimers, we analyzed the crystal structure using the online programs PISA (Protein, Interfaces, Structures, and Assemblies) and EPPIC (Evolutionary Protein-Protein Interface Classifier)(20, 21). These results suggest that AmnC forms a stable symmetric tetramer in the crystal lattice. The tetrameric structure of AmnC was generated by applying crystallographic P6_1_22 symmetry, which is consistent with size-exclusion chromatography data indicating a tetrameric form of AmnC in solution.

**Table 1.** Data collection and refinement statistics.

|  | AmnC |
| --- | --- |
| Wavelength | 0.9798 |
| Resolution range | 25.54 - 1.988 (2.059 - 1.988) |
| Space group | P 61 2 2 |
| Unit cell | a=181.526 b=181.526 c=181.157<br>a=90° b=90° γ= 120° |
| Total reflections |  |
| Unique reflections | 119933 (11656) |
| Multiplicity |  |
| Completeness (%) | 99.74 (98.50) |
| Mean I/sigma(I) | 30.3 |
| Wilson B-factor | 31.54 |
| R<sub>merge</sub><sup>†</sup> | 0.085 (0.839) |
| R<sub>meas</sub> | 0.088 (0.863) |
| R<sub>pim</sub> | 0.020 (0.201) |
| CC1/2 | 0.882 |
| Reflections used in refinement | 119920 (11656) |
| Reflections used for R-free | 6085 (592) |
| R-work | 0.1720 (0.2326) |
| R-free | 0.2078 (0.2704) |
| Number of non-hydrogen atoms | 8355 |
| Macromolecules | 7557 |
| Ligands | 58 |
| Solvent | 740 |
| Protein residues | 983 |
| RMS(bonds) | 0.01 |
| RMS(angles) | 1.22 |
| Ramachandran favored (%) | 96.18 |
| Ramachandran allowed (%) | 3.41 |
| Ramachandran outliers (%) | 0.41 |
| Rotamer outliers (%) | 0.38 |
| Clashscore | 6.21 |
| Average B-factor | 39.11 |
| Macromolecules | 38.31 |
| Ligands | 75.58 |
| Solvent | 44.43 |
| Number of TLS groups | 9 |
| <sup>†</sup> $R_{\text{merge}} = \sum_{hkl} \sum_i \vert I_i(hkl) - \langle I(hkl) \rangle\vert / \sum_{hkl} \sum_i I_i(hkl)$, where $I_i(hkl)$ is an individual intensity measurement and $\langle I(hkl) \rangle$ is the average intensity for all reflections. | |

The overall structure of AmnC protomers shares the general architecture of the ALDH family. The protomeric AmnC is a polypeptide with 491 residues, composed of 16 α-helices, six 3_10_ helices, and 20 β-strands, and comprises three domains: a nicotinamide adenine dinucleotide (NAD+)-binding domain (BD, residues 1–118, 143–253, 447–474), a catalytic domain (CD, residues 254–446), and an oligomerization domain (OD domain, residues 117– 142 and 475–489) (Fig.2). The OD motif is formed by a three-stranded antiparallel β-sheet that is packed in a line and protrudes from the BD domain. The structure of AmnC is highly conserved in the ALDH superfamily. The protein structure comparison server DALI retrieved 49 homologs with similar structures to the AmnC protomer from the protein data bank (PDB) (Table 2)(22). The root-mean-square deviation (RMSD) value estimated based on 262-424 Cα atoms in these structures varied from 0.47–1.15%, with a sequence identity ranging from 36.14–60.79%.

**Figure 2.**
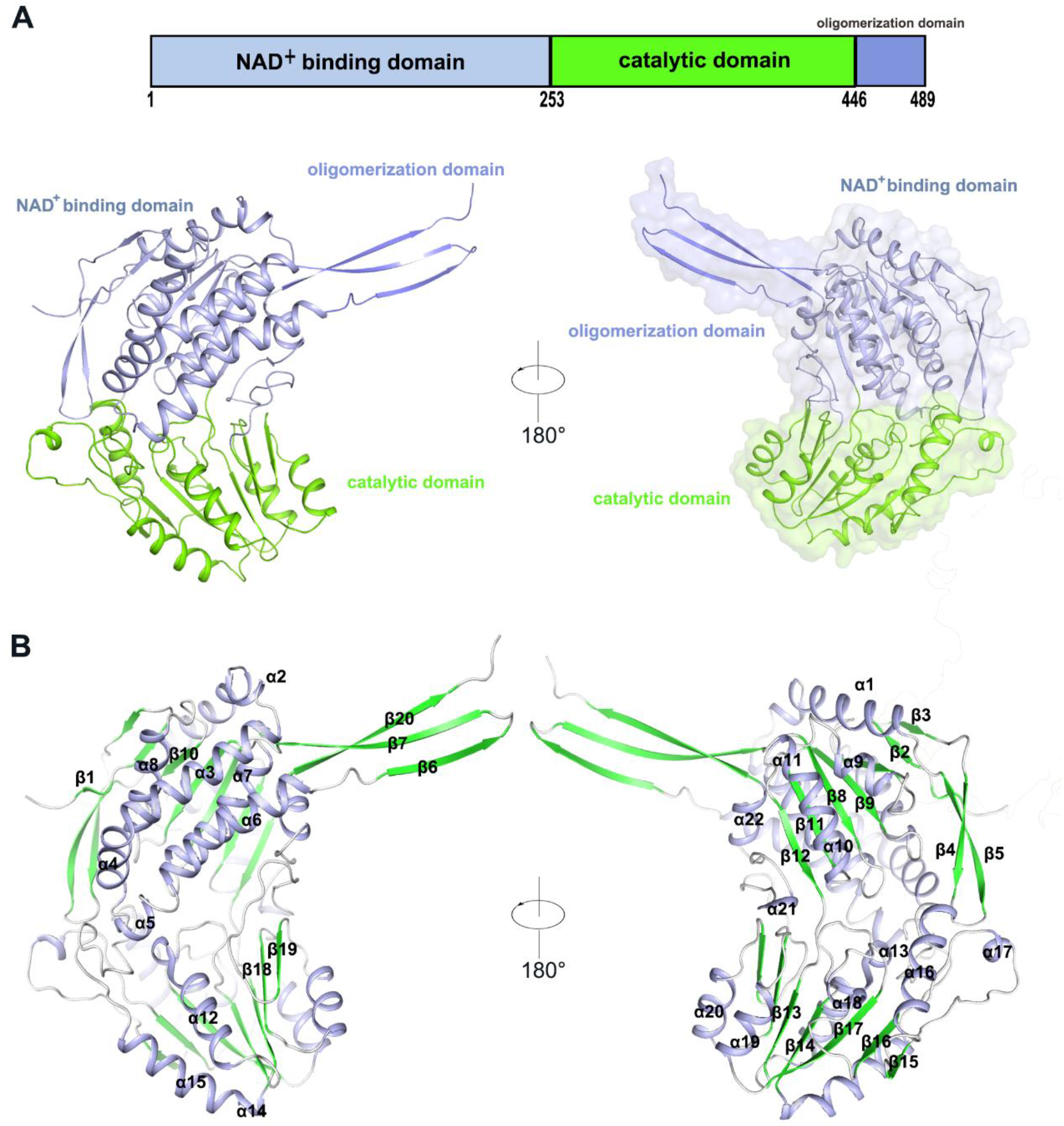
The overall structure of AmnC protomer. (**A**) Domain organization of AmnC, NAD^+^ BD shown in light blue, CD shown in green, and OD shown in slate. Graphical representation of the architecture of AmnC with three domains. Note: the size of the graphic representation does not directly relate to the size of the domains. (**B**) Secondary structure of AmnC promoter. Helixes colored in light blue, β-strands colored in green, and loops colored in gray.

**Table 2.** Homologous proteins of AmnC protomer.

| **PDB code** | **Species** | **C<sub>α</sub> atoms** | **RMSD** | **Sequence identities** |
| --- | --- | --- | --- | --- |
| 4u3w | Burkholderia cenocepacia J2315 | 426 | 0.591 | 63% |
| 3b4w | Mycobacterium tuberculosis H37Rv | 415 | 1.159 | 36% |
| 4fr8 | Homo sapiens | 411 | 0.9 | 37% |
| 2d4e | Thermus thermophilus HB8 | 409 | 0.89 | 40% |
| 2o2p | Rattus norvegicus | 409 | 0.927 | 36% |
| 3iwj | Pisum sativum | 405 | 0.913 | 37% |
| 6pnu | Pseudoalteromonas fuliginea | 405 | 1.442 | 27% |
| 6fk3 | Thermus thermophilus HB27 | 404 | 1.39 | 30% |
| 4nea | Staphylococcus aureus subsp. aureus COL | 399 | 1.125 | 37% |
| 1a4s | Gadus morhua callarias | 397 | 0.99 | 37% |
| 5vbf | Burkholderia vietnamiensis | 396 | 1.409 | 30% |
| 6b4r | Pseudomonas aeruginosa PAO1 | 394 | 0.995 | 36% |
| 4dal | Sinorhizobium meliloti 1021 | 391 | 1.218 | 32% |
| 4ywe | Burkholderia cenocepacia J2315 | 390 | 1.019 | 32% |
| 1t90 | Bacillus subtilis | 390 | 1.162 | 27% |
| 3u4j | Sinorhizobium meliloti | 387 | 0.998 | 34% |
| 4jz6 | Pseudomonas putida | 386 | 1.373 | 28% |
| 5izd | Thermoplasma acidophilum | 380 | 0.91 | 29% |
| 6bjp | Pseudomonas aeruginosa PAO1 | 380 | 1.232 | 38% |
| 3k2w | Pseudoalteromonas atlantica T6c | 377 | 1.141 | 31% |
| 4o5h | Burkholderia cenocepacia J2315 | 376 | 0.861 | 38% |
| 2hg2 | Escherichia coli | 375 | 1.132 | 31% |
| 2bhq | Thermus thermophilus HB8 | 375 | 1.236 | 28% |
| 3ty7 | Staphylococcus aureus subsp. aureus Mu50 | 375 | 1.040 | 33% |
| 4iym | Sinorhizobium meliloti 1021 | 373 | 1.308 | 23% |
| 3rh9 | Marinobacter hydrocarbonoclasticus VT8 | 372 | 1.295 | 32% |
| 2w8p | Homo sapiens | 368 | 1.08 | 35% |
| 3vz2 | Synechococcus sp. PCC 7002 | 368 | 1.265 | 26% |
| 3prl | Alkalihalobacillus halodurans | 368 | 1.456 | 29% |
| 6qhn | Alphaproteobacteria | 368 | 1.701 | 30% |
| 4nmk | Pyrobaculum ferrireducens | 367 | 1.068 | 31% |
| 5j6b | Burkholderia thailandensis E264 | 366 | 1.265 | 28% |
| 3i44 | Bartonella henselae | 365 | 0.96 | 33% |
| 1ky8 | Thermoproteus tenax | 361 | 1.372 | 27% |
| 3ros | Lactobacillus acidophilus | 358 | 1.368 | 26% |
| 2y51 | Paraburkholderia xenovorans LB400 | 357 | 1.708 | 23% |
| 3r64 | Corynebacterium glutamicum ATCC 13032 | 355 | 1.477 | 31% |
| 3efv | Salmonella enterica subsp. enterica serovar Typhimurium | 353 | 1.276 | 26% |
| 6rts | Streptomyces clavuligerus | 351 | 1.185 | 28% |
| 4e3x | Mus musculus | 339 | 1.545 | 24% |
| 5u0l | Marinobacter hydrocarbonoclasticus VT8 | 338 | 1.14 | 27% |
| 5mz5 | Physcomitrium patens | 337 | 1.163 | 29% |
| 4dng | Bacillus subtilis | 337 | 1.249 | 25% |
| 6k0z | Staphylococcus aureus | 318 | 1.877 | 21% |
| 4h7n | Trichormus variabilis ATCC 29413 | 311 | 1.41 | 28% |
| 4o6r | Burkholderia cenocepacia | 297 | 0.92 | 38% |

Six ALDH superfamily protein structures were subsequently chosen to represent all prokaryotic and eukaryotic proteins in this protein family. These include the *Pseudomonas aeruginosa* aldehyde dehydrogenase PA5312 (PDB code:6B4R), *Mycobacterium tuberculosis* aldehyde dehydrogenase family protein MT0233 (PDB code: 3B4W), the *moss* predicted protein PHYPA_001937 (PDB code:5MZ5), *Pisum sativum* aminoaldehyde dehydrogenases (AMADHs) (PDB code:3IWJ), *Rattus norvegicus* 10-formyltetrahydrofolate dehydrogenase (Ct-FDH) (PDB code:2O2P), and *Homo sapiens* succinic semialdehyde dehydrogenase (SSADH) (PDB code:2W8P)(23-26). Pairwise superposition of the six protomers yielded RMSD values equal to 1.02 Å (for 384 Cα atoms, AmnC/ PA5312), 1.16 Å (for 415 Cα atoms, AmnC/ MT0233), 1.16 Å (for 337 Cα atoms, AmnC/ PHYPA_001937), 0.91 Å (for 307 Cα atoms, AmnC/ AMADHs), 0.92 Å (for 409 Cα atoms, AmnC/ Ct-FDH), and 1.08 Å (for 368 Cα atoms, AmnC/ SSADH) (Fig.3).

**Figure 3.**
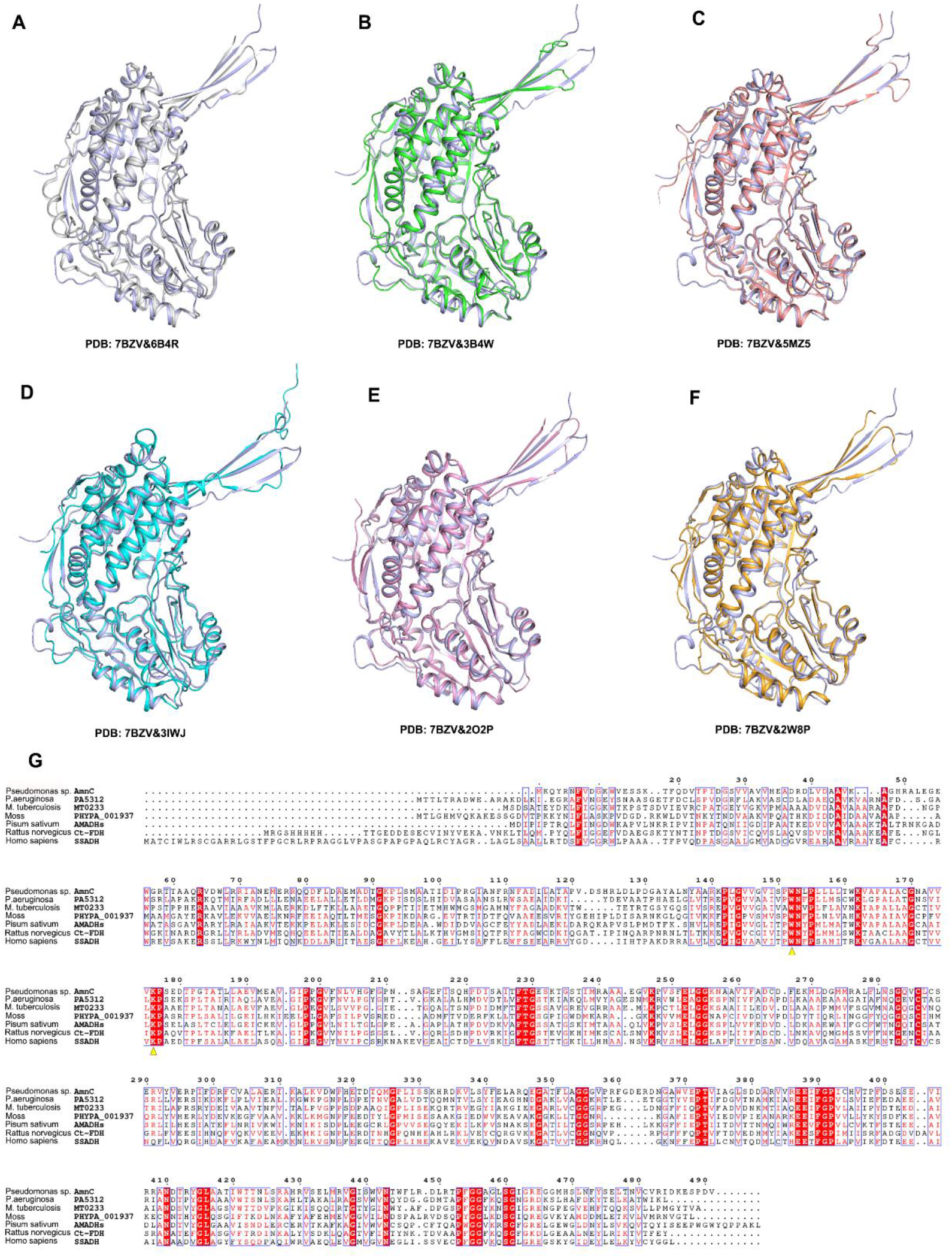
Structural superimposition and multiple sequence alignment of AmnC with ALDH family members. (**A–F**) Superposition of AmnC protomer from *Pseudomonas* sp. *AP–3* (light blue) with the protomeric structures of PA5312 from *P. aeruginosa* (RMSD = 1.02 Å), MT0233 from *M. tuberculosis* (RMSD = 1.16 Å), PHYPA_001937 from *Moss* (RMSD = 1.16 Å), AMADHs from *Pisum sativum* (RMSD = 0.91 Å), Ct-FDH from *Rattus norvegicus* (RMSD =0.92 Å), and SSADH from *homo sapiens* (RMSD = 0.91 Å). (**G**) Sequence alignment of AmnC with other members of the ALDH family. Residues labeled with yellow triangles are conserved residues associated with the NAD^+^ binding pocket.

### The tetrameric assembly of AmnC

The quaternary structure of AmnC shows that AmnC assembles into a tetramer consisting of a dimer of dimers. Two protomers assemble into a solid dimer, and then two dimers assemble to form a tetramer (Fig.4). First, two AmnCs form a head-to-tail dimer. Each protomer accessible surface buried in the dimer is 2641.4 Å^2^, and the proportion of the buried area to the protomeric total surface area is approximately 50%. The major dimer interface consists of the OD (β6, β7, and β20) from one of the protomers stretching into the CD of the opposite protomer. Meanwhile, a pair of antiparallel ⟨−ηελιχεσ in the dimer composed of helix α10 and extending to strand β12 from each subunit of the dimer acts as a minor dimer interface. Hydrophobic interactions are a major contribution to stabilizing the dimer interfaces, which tend to form more hydrogen bonds and salt bridges (Fig.5*A*). Then, the tetramer interface is generated by residues V121 and T423 from one of the AmnC dimers forming hydrogen bonds with residues S123 and L425 in the dimeric partner (Fig.5*B*). To complete the tetramer, two β6 strands in the OD from one of the dimers contact the corresponding two β6 strands of the neighboring dimer to complete two β-barrel-like folds that span four subunits. To further identify the amino acid residues required for tetramer or dimer interfaces, we prepared two mutants based on the AmnC tetramer structure. The mutants, AmnC ΔOD1 (Δ477-491 and Δ124-138) and AmnC ΔOD2 (S123G, H124G, L425A), are expected to be unable to form tetramers from dimers (Fig.6*A, B*). The oligomerization states of the AmnC mutants were evaluated by size-exclusion chromatography (SEC) and dynamic light scattering (DLS). The truncated mutants AmnC ΔOD1 (Δ477-491 and Δ124-138) and AmnC ΔOD2 (S123G, H124G, L425A) exhibited larger retention volumes than AmnC WT; AmnC ΔOD1 was eluted at 14.04 mL on a Superdex 200 size-exclusion column, while AmnC ΔOD2 was eluted at 14.4 mL. The molecular mass of the mutants was 73.67kD and 67.45kD for AmnC ΔOD1 and AmnC ΔOD2, respectively, approximately corresponding to the mass of a dimer (Fig.6*C, D*). DLS analysis was performed to assess the multimeric state of each protein. AmnC WT appears to be predominantly tetrameric, with an estimated molecular mass of 222 kDa (Z-average size of 15.52 d.nm and index of polydispersity PdI of 0.11). AmnC ΔOD1 and AmnC ΔOD2 were calculated to have an Z-average size of 7.52 d.nm (PdI of 0.21) and 7.90 d.nm (PdI of 0.17), with estimated molecular masses of 74.70 kDa and 83.80 kDa, respectively, and appear to be predominantly dimeric (Fig.6*E, F*).

**Figure 4.**
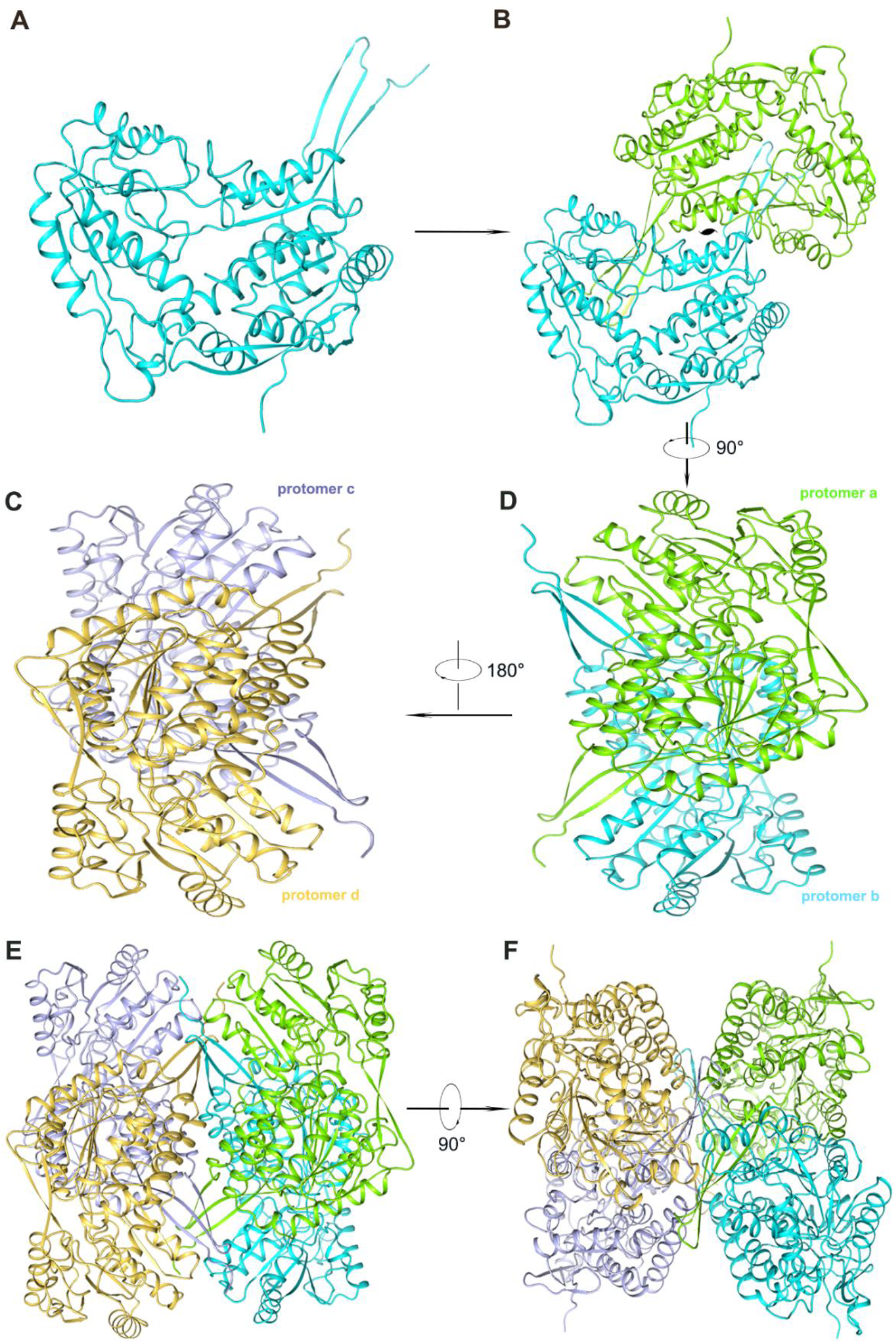
Overview of the tetrameric assembly of AmnE. (**A**) Three-dimensional view of the AmnC protomer. (**B**) AmnC dimer. Subunits are drawn in different colors in this ribbon diagram viewed along the two-fold axis. (**C–F**) The AmnC tetramer is composed of two identical dimers (I and II). Dimer I consists of protomer a and b, dimer II consists of protomer c and d.

**Figure 5.**
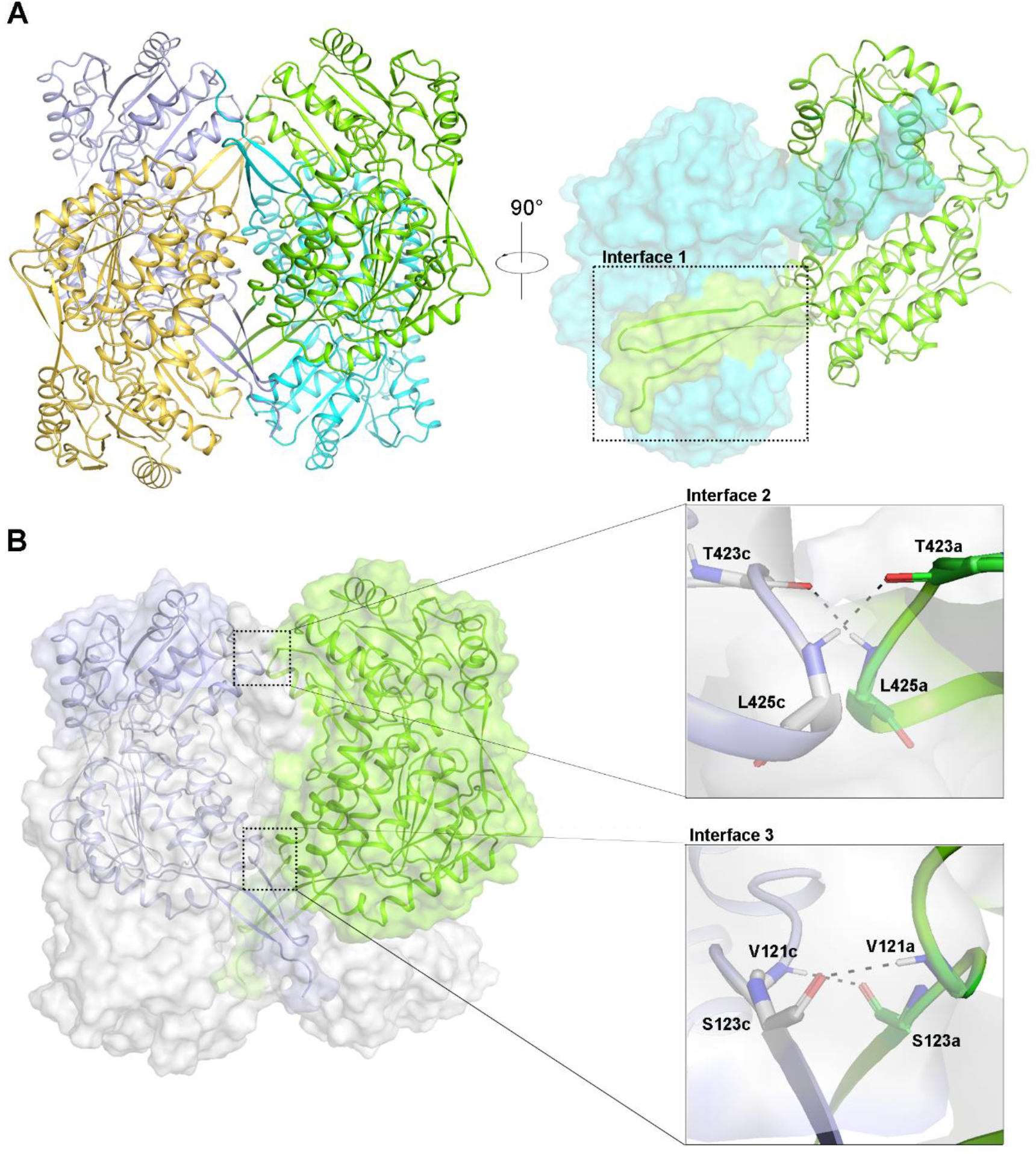
AmnC tetrameric interface. (**A**) Interface 1 shows that the OD stretches into the CD. (**B**) T423 forms hydrogen bonds with L425 from the CD of its dimer partner shown in interface 2. Interface 3 is provided by V121 forming hydrogen bonds with S123 in the OD.

**Figure 6.**
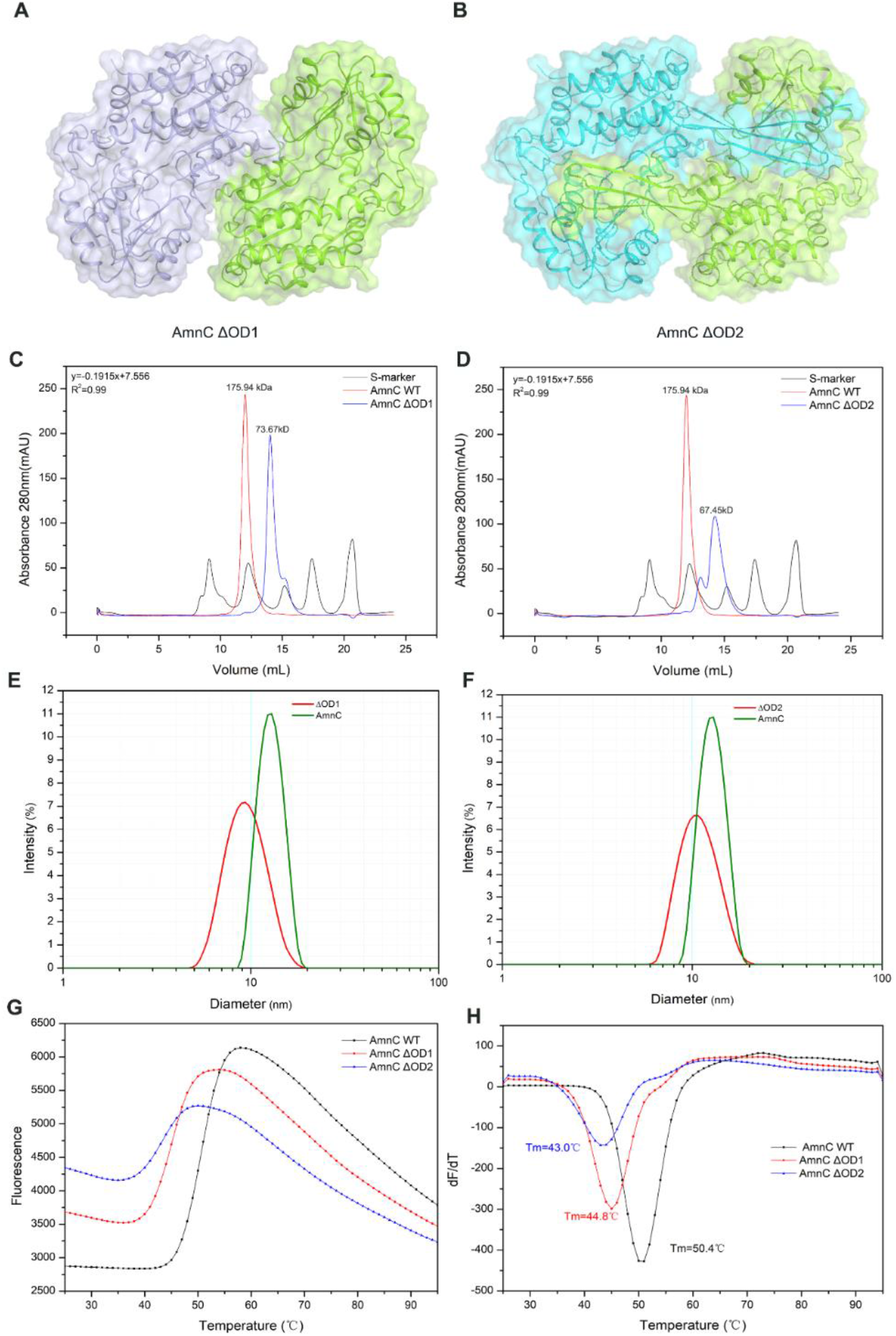
Biochemical characterization of AmnC tetramerization. (**A**) Schematic of the AmnC mutant AmnC ΔOD1. (**B**) Schematic of the AmnC mutant AmnC ΔOD2. (**C-D**) Analytic SEC of AmnC-WT, AmnC ΔOD1, and AmnC ΔOD2. The red solid curve refers to AmnC-WT, the blue solid curve refers to AmnC ΔOD1 or AmnC ΔOD2, and the black solid curve refers to the molecular weight standards. (**E-F**) DLS analysis of AmnC-WT, AmnC ΔOD,1 and AmnC ΔOD2. The green solid curve refers to AmnC-WT, the red solid curve refers to AmnC ΔOD1 or AmnC ΔOD2. (**G**) DSF recording of fluorescence intensity versus temperature for AmnC-WT, AmnC ΔOD1, and AmnC ΔOD2. (**H**) First derivatives versus temperature for AmnC-WT, AmnC ΔOD1, and AmnC ΔOD2.

To further assess the protein folding state and thermal stability, we conducted differential scanning fluorimetry (DSF) experiments. Based on the change in fluorescence intensity as the temperature was gradually increased, we found that the melting temperature (Tm) value of AmnC-WT was 50.4°C, 44.8 °C for AmnC ΔOD1, and 42.1°C for AmnC ΔOD2 (Fig.6*G, H*). The dimeric forms of AmnC have lower melting temperatures compared to AmnC-WT, indicating that the AmnC tetramer is much more stable than the AmnC dimers. The small changes in Tm values (44.8 °C, 42.1°C) between the AmnC variants also indicated that AmnC ΔOD2 preserves the high-energy fold relative to ΔOD1.

### The tetrameric structure of AmnC in solution

PDBePISA was used to analyze AmnC crystal packing and predicated two stable assemblies, including a head-to-tail dimer (R_g_ = 30.1 Å) and the tetramer (R_g_ =38.4 Å). Considering the mutants ΔOD1 (R_g_ = 29.83 Å) and ΔOD2 (R_g_ =30.12 Å) with diverse dimeric forms, all potential oligomers are depicted in Figure 7A. The quaternary structure of AmnC and its variants in solution was studied using SAXS. AmnC ΔOD2 shows a high aggregation tendency under various conditions and was not used for further measurements. SAXS curves were collected from AmnC-WT and AmnC ΔOD1 at different protein concentrations (Table 3). Qualitatively, the shape of the scattering curve from AmnC-WT remains constant with increasing protein concentration and consistently displays a small bump near *q* =0.075–0.15 Å^−1^. Guinier plots yielded a radius of gyration (Rg) in the range of 38–39 Å, and, given that the Rg of AmnC-WT calculated from the crystal structure is 38.4 Å, this suggests that AmnC forms a tetramer in solution (Fig.7*B*). While the SAXS curves from AmnC ΔOD1 show a small bump near *q* =0.1–0.175 Å^−1^, the Guinier plots yield a Rg in the range of 28–29 Å (Fig.7*C*). Kratky plots demonstrate that the SAXS data are predominantly from folded particles. The peak for AmnC ΔOD1 was slightly shifted to the right, as expected for its slightly elongated shape, and the small increase evident at qRg > 1 suggests some flexibility (Fig.*7D*). The calculated distance distribution function suggests that the maximum particle dimension (Dmax) of AmnC ΔOD1 was 88–93 Å, which is much smaller than that of AmnC WT at 115–121 Å (Fig.*7E*). To further characterize the tetrameric state of AmnC, AmnC was eluted at 12.69 mL on a Superdex 200 10/300GL size-exclusion column with a calculated molecular mass of 175.94 kDa (the theoretical monomer mass is 53.72 kDa), which is close to that AmnC as a tetramer (Fig.6*C*). Theoretical SAXS scattering curves were generated based on models derived from crystal structures for comparison with experimental data using CRYSOL(27). The theoretical SAXS scattering curves generated from atomic coordinates obtained from the crystal structure of the AmnC tetramer faithfully reproduced the experimental data, and the consistency of the AmnC ΔOD1 dimeric model with the experimental data was also reproduced (Fig.7*F, G*). SAXS modeling of AmnC and AmnC ΔOD1 was performed using DAMMIN(28). The atomic model was docked into an ab initio envelope using the program SUPCOMB(29). We observed that the AmnC tetrameric crystal structure and AmnC ΔOD1 dimeric models match the SAXS model in three ways. These results suggest that AmnC appears to exist as a tetramer in solution and AmnC ΔOD1 is a dimer in solution.

**Figure 7.**
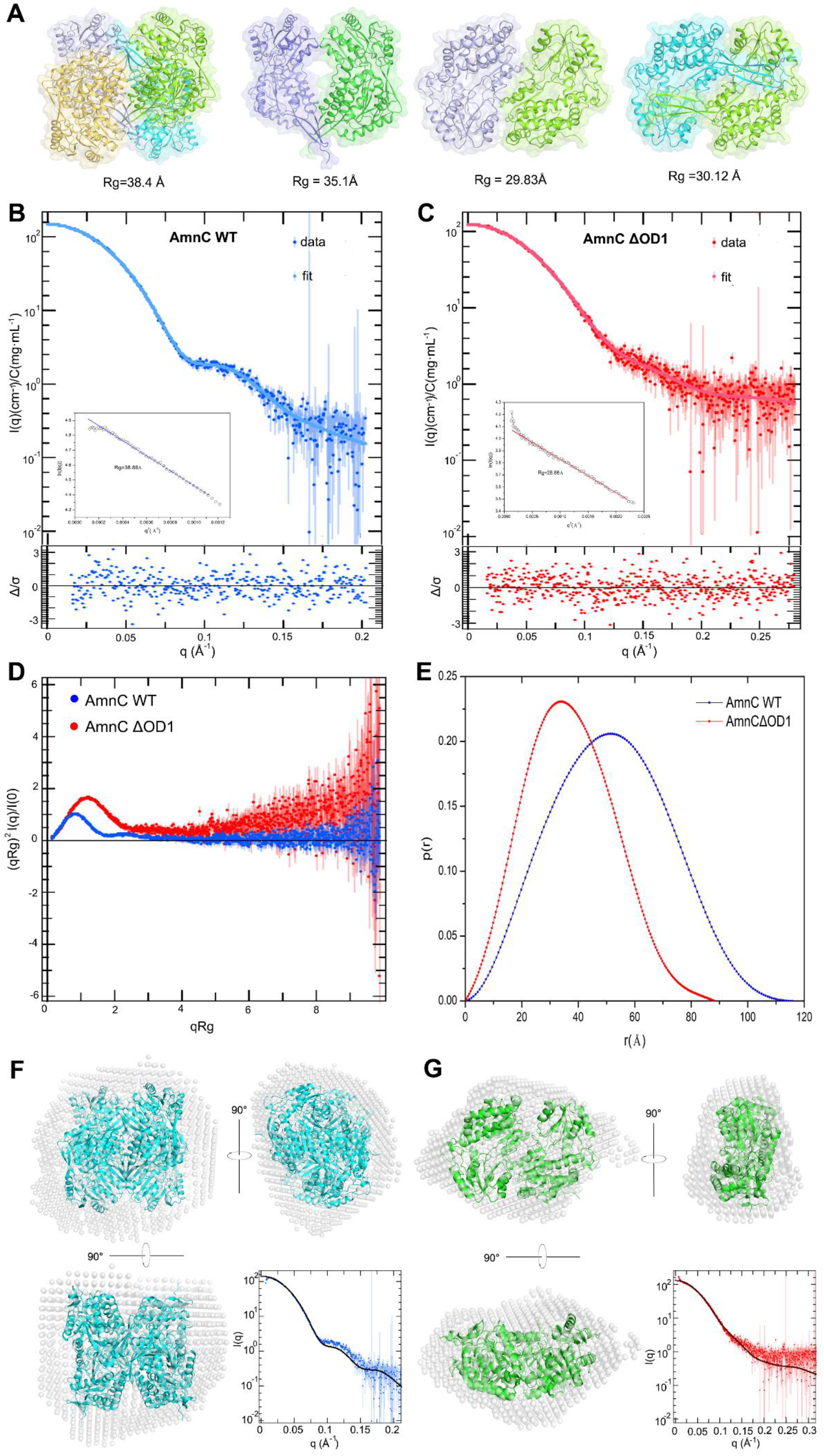
SAXS analysis of AmnC and AmnC ΔOD1. (**A**) Potential oligomers of AmnC WT tetrameric and dimeric form. The Rg value calculated by using CRYSOL, shown below. (**B**) SAXS curves for AmnC collected at a protein concentration of 1.0 mg/mL. The lower left quarter inset shows Guinier plots. The lower inset plot is the error-weighted residual difference plot Δ/σ = [*I*_exp_(*q*) - *cI*mod(*q*)]/ σ(*q*) *versus q*. (**C**) SAXS curves for AmnC ΔOD1 collected at a protein concentration of 1.0 mg/mL. The lower left quarter inset shows Guinier plots. The lower inset plot is the error-weighted residual difference plot Δ/σ = [*I*_exp_(*q*) - *cI*mod(*q*)]/ σ(*q*) *versus q*. (**D**) Kratky plots transformed from the SAXS profiles. (**E**) The pairwise distance distributions derived from the SAXS profiles using GNOM within the ATSAS software suite. (**F**) AmnC-WT DAMMIN model (gray spheres) overlaid with the crystal structure. Model overlays were optimized using SUPCOMB and are shown in three views. The model fits to *I(q)* versus *q* for AmnC-WT at the lower right corner. (**G**) AmnC ΔOD1 DAMMIN model (gray spheres) overlaid with the crystal structure. Model overlays were optimized using SUPCOMB and are shown in three views. The model fits to *I(q)* versus *q* for AmnC ΔOD1 shown in the lower right corner.

**Table 3.** SAXS parameters of AmnC WT and AmnCΔOD1

| Parameters |  |  |  |  |
| --- | --- | --- | --- | --- |
| Data Collection |  |  |  |  |
| Beamline name | SSRF (Shanghai, China) BL19U2 |  |  |  |
| Source type | Undulator, U20 |  |  |  |
| Mirrors | 1.0m-long Rh-coated focusing mirror,3.5 mrad |  |  |  |
| Energy range(keV) | 7-15 |  |  |  |
| Wavelength range (Å) | 0.82-1.77 |  |  |  |
| Beam size (detector plane) | 0.33mm(H) ×0.05mm(V) |  |  |  |
| Flux (photons s<sup>-1</sup>) | 5×10<sup>12</sup>(at 1.03 Å) |  |  |  |
| Sample loading | Robot |  |  |  |
| Cryo capability (K) | 277-333 |  |  |  |
| Sample cell unit | Quartz capillary |  |  |  |
| Detector type | CMOS hybrid pixel |  |  |  |
| Detector model | Pilatus 1M |  |  |  |
| *q* range(nm<sup>-1</sup>) | 0.04-4.5(at 1.03 Å) |  |  |  |
| Structural Parameters |  |  |  |  |
|  | AmnC WT |  | AmnC ΔOD1 |  |
|  | 1mg/mL | 3mg/mL | 1mg/mL | 5mg/mL |
| Guinier analysis |  |  |  |  |
| Rg(Å) | 38.86±0.13 | 39.63±0.06 | 28.86±0.14 | 29.01±0.06 |
| q<sub>min</sub>(Å<sup>-1</sup>) | 0.01067 | 0.01067 | 0.01209 | 0.00782 |
| qRg range | 0.41-1.30 | 0.42-1.26 | 0.35-1.35 | 0.23-1.28 |
| P(r) analysis<sup>a</sup> |  |  |  |  |
| Rg(Å) | 39.21 | 39.44 | 28.44 | 28.78 |
| D<sub>max</sub>(Å) | 115.85 | 121.03 | 88.55 | 93.21 |
| q-range (Å<sup>-1</sup>) | 0.0107-0.0334 | 0.01067-0.03201 | 0.0121-0.0467 | 0.0078-0.0443 |
| Total estimate | 0.99 | 0.93 | 0.92 | 0.95 |
| Porodvolume(Å<sup>3</sup>) | 326328 | 328332 | 116779 | 129303 |
| Modeling |  |  |  |  |
| DAMMIN χ<sup>2</sup> |  | 1.033 | 0.907 |  |
| DAMMIN Ensemble Resolution |  | 38±3 | 28±2 |  |
| DAMMIN NSD |  | 0.556±0.01 | 0.541±0.02 |  |
| Software Employed |  |  |  |  |
| Primary data Processing | RAW |  |  |  |
| *P*(*r*) | GNOM |  |  |  |
| *Ab initio* shape analysis | DAMMIF |  |  |  |
| Rigid body modeling | CORAL |  |  |  |
| SAXS Profile computation | CRY SOL |  |  |  |
| Molecular Visualization | PyMOL |  |  |  |
| <sup>a</sup> From calculations of the distance distribution function using GNOM via PRIMUS. |  |  |  |  |

### Tetrameric assembly is required for AmnC catalytic activity

Upon testing the catalytic activity of AmnC ΔOD1, we found that there was no reduction in absorbance when AmnC ΔOD1 was incubated with NAD^+^ and 2-HMS, and the absorption maximum at 375 nm did not decrease when compared with AmnC-WT (Fig.8*D, E*). Experimental data showed that AmnC ΔOD1 lost dehydrogenase activity. Superposition AmnC with *Homo sapiens* ALDH1A3 indicated that the NAD^+^ binding pocket is highly conserved (Fig.8*A*). Residues K176 and W152 interact with NAD^+^ and are conserved, as verified by sequence alignment with the ALDH superfamily (Fig.3*G*). We measured the affinity of AmnC-WT and AmnC ΔOD1 for NAD^+^ and found that the affinity of AmnC-WT was 63±24 µM while that of AmnC ΔOD1 was only 3±0.3mM (Fig.8*B, C*), indicating that AmnC ΔOD1 loses the ability to bind with the cofactor NAD^+^, potentially causing catalytic inactivity. These data emphasize the importance of AmnC oligomerization for enzymatic activity. AmnC is unique in that its enzymatic activity is oligomeric state–dependent, with tetramerization being required for its function.

**Figure 8.**
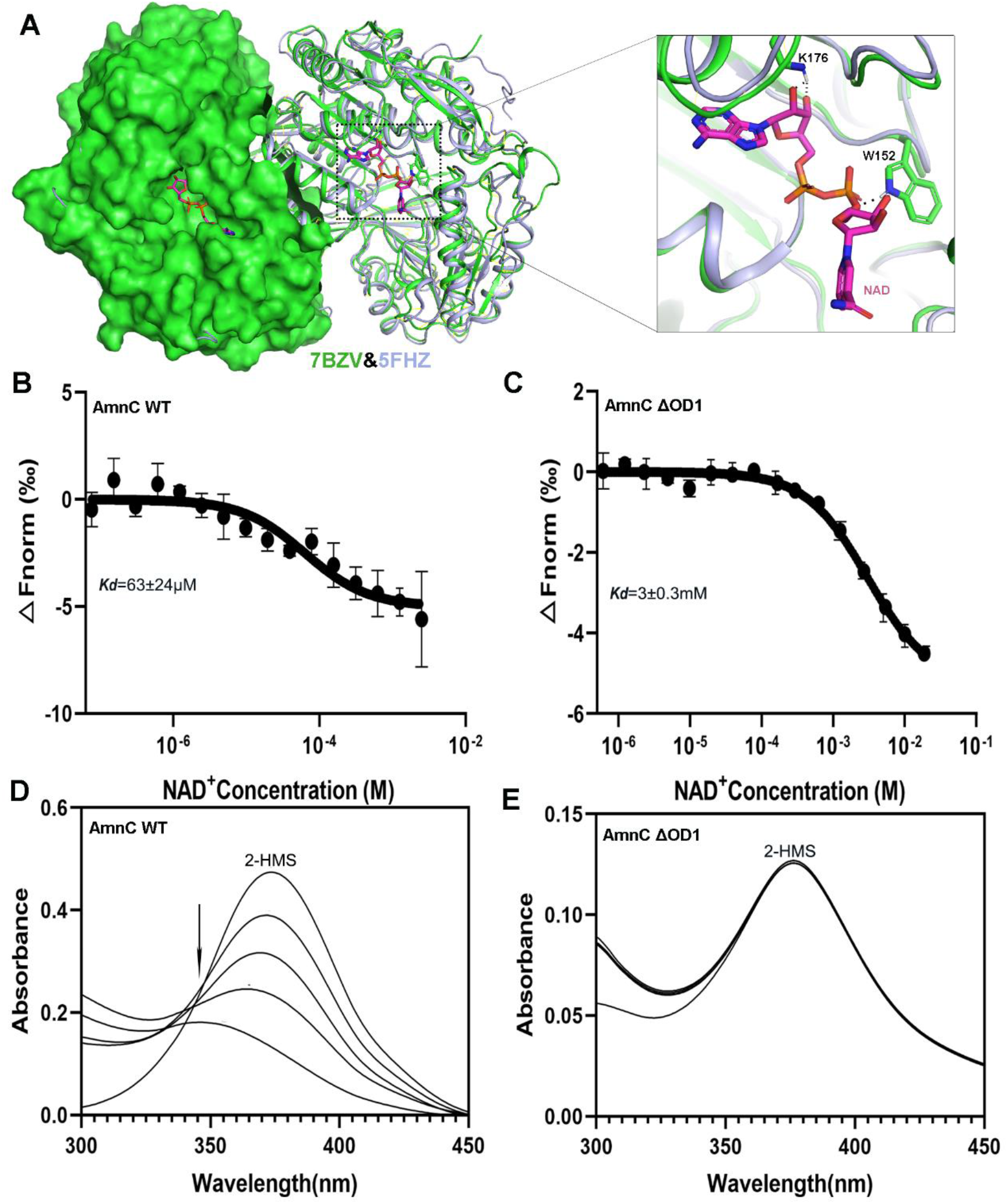
Catalytic activity of AmnC ΔOD1. (**A**) Superposition of AmnC (PDB:7BZV) with ALDH1A3(PDB:5FHZ) from *Homo sapiens* shows that the NAD^+^ binding pocket is conserved, and the conserved residues shown as sticks. (**B**) Dose-response curve for the binding interaction between AmnC and NAD^+^; the binding curve yields a Kd of 63 µM. (**C**) Dose-response curve for the binding interaction between AmnC ΔOD1 and NAD^+^; the binding curve yields a Kd of 3 mM. (**D**) Representative assay showing the activity of AmnC on 2-HMS (λmax 375 nm). (**E**) Representative assay showing the activity of AmnC ΔOD1 on 2-HMS (λmax 375 nm).

## Discussion

2-aminophenol compounds are intermediates in the biodegradation of nitrobenzene, an extremely hazardous substance released into the environment by anthropogenic activities. The metabolic pathway of 2-aminophenol in *Pseudomonas* sp. *AP-3* is known as the meta-cleavage pathway, in which 2-aminomuconic 6-semialdehyde is converted to 2-aminomuconic acid by AmnC. In this study, we elucidated the structure of AmnC and how it is involved in the meta-cleavage pathway in *Pseudomonas* sp. *AP–3*. AmnC belongs to the ALDH superfamily, members of which catalyze the NAD^+^-dependent oxidation of aldehydes to carboxylic acids and are present in all domains of life.

From a structural view, ALDH enzymes typically exist and function as homotetramers or homodimers; meanwhile, a small number of structures have demonstrated the presence of hexameric enzymes. AmnC has a dimer of dimer quaternary structure in which the dimer interface is primarily composed of beta strands β19 and β6 from the OD (residues 118-142 and 474-191) and is made up of relatively stronger interactions than the tetramer interface associated with residues V121, S123, T423, and L425. Mutagenesis and biochemical studies have shown that hydrogen bonding and hydrophobic interactions stabilize the dimer and tetramer interfaces, and ionic interactions in the NAD^+^ BD also play pivotal roles in the dimer interface organization. Our data clearly indicate that the stability of the AmnC tetramer is predominantly affected by tetramer interfaces, despite that the buried area of the dimer interface is 2,637 Å^2^, which is three times larger than that of the tetramer interface (714 Å^2^). Disruption of the tetramer interface destabilizes the enzyme. The first evidence supporting this hypothesis is the observation that during the thermal stability experiments, the Tm of AmnC-WT was 6-7°C higher than that of the tetramer interface mutant AmnC ΔOD2 (S123G, H124G, L425A), indicating that the dissociation of subunits at the tetramer interface significantly affects the stability of AmnC. Second, the mutant AmnC ΔOD2 is highly unstable during the recombinant protein purification process, posing a challenge for long-term storage and further testing. However, the dimer interface mutant AmnC ΔOD1 (Δ477-491 and Δ124-138) was more stable than the tetramer interface mutant ΔOD2. Gel filtration chromatography and SAXS data indicated that AmnC ΔOD1 also formed a dimer in solution. Because the four subunits have identical structural characteristics, the difference in structural stability between the two mutants can be largely attributed to the differences in subunit-subunit interactions between the tetramer and dimer interfaces of AmnC.

Our data also indicate that the enzyme activity of AmnC is affected by disruption at the dimer interface. There were significant differences in the enzyme activity between AmnC-WT and AmnC ΔOD1. The mutant completely lost catalytic activity, suggesting that the four subunits of AmnC interact with one another and function cooperatively. We also attempted to prepare a monomeric form of AmnC by multiple mutations at the dimer and tetramer interfaces; however, this approach was successful due to aggregation of the monomers. Given these results, we propose that the functional significance of the tetrameric structure of AmnC may be in maintaining the stability of the enzyme. Meanwhile, the mutant ΔOD1 lost the ability to bind the cofactor NAD^+^, which may indicate that the quaternary structure of AmnC is required for NAD^+^ binding.

In summary, we demonstrated that the tetrameric form of AmnC is required for proper enzyme activity. The quaternary structure of AmnC is primarily responsible for protein stability. Dissociation of subunits at the dimer or tetramer interface disrupts protein stability. In the dimer interface mutant, the ability to bind the cofactor NAD^+^ is lost, causing the enzyme to be inactive. Thus, the tetramer organization of AmnC is principally responsible for enzyme activity.

## Materials and Methods

### Generation of wild-type and mutant plasmid constructs

The gene encoding AmnC in *Pseudomonas* sp. *AP–3* was synthesized by Sangon Biotech Co., Ltd., China, and subcloned into the bacterial vector pET–28a (GE Healthcare, United States) while introducing six histidine residues (6×His-tag) at the N-terminus of the gene. To analyze the oligomerization interfaces in AmnC, we generated two mutants, AmnC ΔOD1 (Δ124-138 and Δ477-491) and AmnC ΔOD2 (S123G, H124G, and L425A). The fragment of AmnC ΔOD1 was inserted into pET-28s (GE Healthcare, United States) with a 6×His-SUMO tag at the N terminus, and AmnC ΔOD2 was inserted into pET-28a using the FastCloning method. All plasmids were sequenced to confirm their identity using commercial DNA sequencing (Sangon Biotech, China). All primers used in this study are listed in Table 4.

**Table 4.**
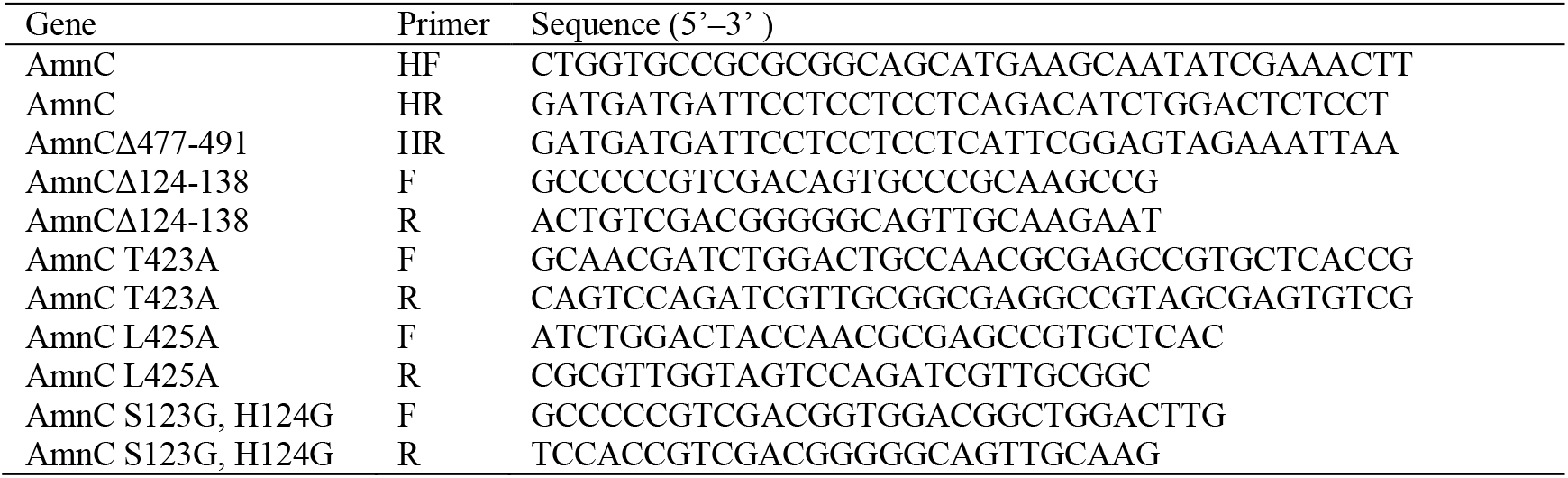
Primers used in this study.

### Recombinant protein expression and purification

Wild-type and mutant AmnC were expressed in *Escherichia coli* BL21 (DE3) (TransGen Biotech) grown on Luria-Bertani (LB) medium containing 50 mg/ml kanamycin at 37 °C. When the optical density (OD600) value was 0.6–0.8, recombinant protein expression was induced at 16°C for 12 h with 0.2 mM isopropyl-b-D-1-thiogalactopyranoside (IPTG). Cells were harvested by centrifugation for 10 min at 4,000 rpm, resuspended in lysis buffer (20 mM Tris–HCl, 200 mM NaCl, and 10 mM imidazole, pH 8.0), and lysed using a high-pressure homogenizer (JNBIO, China) at 6°C. Cellular debris was removed by centrifugation for 30 min at 15,000 rpm at 4 °C (Thermo Sorvall LYNX 6000, United States). The supernatant was loaded into a 5 mL Ni–NTA affinity chromatography column (GE Healthcare, United States) that was pre-equilibrated with lysis buffer. The bound protein was eluted with elution buffer (20 mM Tris–HCl, 200 mM NaCl, 250 mM imidazole, pH 8.0). Thrombin (Sigma, United States) and SUMO protease (Ulp1) were used to cleave the N-terminal 6×His tag and 6×His-SUMO tag, respectively. Protein samples were loaded into dialysis tubing with buffer (20 mM Tris–HCl, 20 mM NaCl, 5% (v/v) glycerol, pH 8.0) and dialyzed at 16 °C overnight.

After the removal of epitope tags, recombinant proteins were further purified by anion-exchange chromatography on a Resource Q column (GE Healthcare, United States) that was pre-equilibrated with buffer A (20 mM Tris–HCl, 20 mM NaCl, pH 8.0) and eluted with buffer B (20 mM Tris–HCl, 1M NaCl, pH 8.0) using a linear gradient. The samples were further purified by gel filtration chromatography on a Superdex200 10/300 GL column (GE Healthcare, USA) equilibrated with 20mM Tris–HCl, 150 mM NaCl, 1mM DTT, 1mM EDTA, 5% (v/v) glycerol, pH 8.0. All proteins were concentrated by ultrafiltration with Amicon Ultra (10 kDa, Millipore, Germany) and protein purity was confirmed by SDS– PAGE, followed by staining with Coomassie Brilliant Blue R-250. For crystallization purposes, wild-type AmnC was concentrated to 10 mg/mL.

### Protein crystallization, X-ray diffraction data collection, and structure determination

Initial crystallization trials were performed using a Gryphon robot (Art Robbins Instruments, United States) using the sitting-drop vapor diffusion method at 20°C. Drops were prepared by mixing 0.3 μL purified protein and reservoir solution. This screening consisted of using commercial crystallization kits (Crystal Screen, Crystal Screen 2, Index Screen, Salt screen, and PEG Screen) purchased from Hampton Research (United States). Protein crystals were obtained in 1.8 M ammonium sulfate, 0.1 M BIS-TRIS pH 6.5, 2% v/v polyethylene glycol monomethyl ether 550. Optimization was carried out in 24-well plates using the hanging-drop method. Drops were mixed with 1 μL of protein and 1 μL of reservoir solution. A high-quality crystal was obtained in 1.8 ammonium sulfate, 0.1 M BIS-TRIS pH 6.5, 4.4% v/v polyethylene glycol monomethyl ether 550.The crystals were flash-cooled in liquid nitrogen and diffraction data were collected at the Shanghai Synchrotron Radiation Facility (SSRF) using a beamline BL19U1 (Shanghai, China). Data were processed using HKL2000 software(31). The best crystal diffracted X-rays formed a pattern with a resolution of 1.9 Å. The space group of AmnC is P6_1_22, with cell dimensions set at a = 181.5 Å, b = 181.5 Å, c = 181.2 Å. The AmnC structure was solved by the molecular replacement method with the Phaser in the CCP4 suite(32), using the initial model of *Pseudomonas fluorescens* AMSDH (PDB: 4NPI). The refined model was further constructed manually according to the electron density map with COOT, and the quality of the refined models was assessed using PHENIX(33, 34). The final structure was derived to a R factor of 17.2% (R_free_ factor of 20.78%). The data collection, statistics, and structure refinement statistical outcomes are listed in Table 2.

### Small-angle X-ray scattering

SAXS data were collected using a beamline BL19U2 at the SSRF (Shanghai, China). The beamline details are listed in Table 3. AmnC was purified and verified using SEC. Serially diluted AmnC samples (1–5 mg mL^−1^) were collected. All samples were prepared in 20 mM HEPES, 150 mM NaCl, pH 7.0. Each sample (60 µL) was continuously passed through a capillary tube exposed to a 240 × 50 μm X-ray beam. For each protein, 20 images were collected at three different protein concentrations. Images for background subtraction were collected in a similar manner to dilute the buffer.

All SAXS data were processed using the BioXTAS RAW (35) and ATSAS 2.7 software packages. The scattering profile of the protein was obtained by subtracting the buffer profile. The forward scattering, I(0), and the radius of gyration, Rg, were evaluated using the Guinier approximation. The maximum dimension, Dmax, and interatomic distance distribution function (P(r)) were calculated using the program GNOM(36). Ab initio modeling of the molecular envelope was calculated and optimized using the programs DAMMIF (37) and DAMMIN (28). The theoretical scattering curve from the atomic model was calculated and compared with the experimental curve obtained by CRYSOL(38). The atomic model was docked into an ab initio envelope using the program SUPCOMB (39).

### Analytic size-exclusion chromatography

Analytic SEC was adopted to investigate the oligomeric state of AmnC and its variants using a Superdex 200 10/300 GL column (GE Healthcare, United States). The column was pre-equilibrated with analysis buffer (50 mM Tris-Cl, 150 mM NaCl, 1 mM DTT, 1 mM EDTA, and 5% (v/v) glycerol, pH 8.0). Furthermore, all samples were prepared at a concentration of 5 mg/mL and analyzed at a flow rate of 0.3 mL/min at 16°C. The molecular mass was calculated using the following equation:.

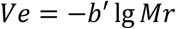

*Ve* is the volume at which the intermediate molecules elute, and *Mr* is the molecular mass.

### Dynamic light scattering

The hydrodynamic radius and polydispersity of the purified samples were estimated using a DLS assay performed on a Zetasizer mV (Malvern Instruments, Worcestershire, UK). All samples were centrifuged for 45 min at 50,000 rpm at 4 °C prior to DLS. The final protein concentration for DLS measurements was approximately 1 mg/mL in 20 mM HEPES, 150 mM NaCl, 5%(v/v) glycerol, pH 7.5 (200 μL, 16 °C).

### Differential scanning fluorimetry

DSF was adopted to obtain the temperature midpoint for the protein unfolding transition Tm. Solutions of 0.1 μL 5000×Sypro orange dye (Sigma) and 100 μL 5μM proteins were mixed together, and 10μL of the mixture was added to the wells of a 96-well thin-wall PCR plate (Bio-Rad). The plates were heated in an iCycler iQ Real Time PCR Detection System (Bio-Rad) from 25 to 95°C in increments of 1°C/30 s. Each protein sample was analyzed in triplicate.

### Microscale thermophoresis

Proteins were labeled using the Monolith NT Protein Labeling Kit RED (NanoTemper Technologies) according to the manufacturer’s protocol and using a buffer consisting of 20 mM HEPES, pH 7.2, 150 mM NaCl, and 0.05% Tween-20. The concentration of the labeled proteins was 20 nM. The corresponding unlabeled binding partner was titrated in 1:1 (v/v) dilutions, with the highest final concentration chosen approximately 20-fold above the Kd value. Thus, the highest final concentration was 62.5 μM NAD^+^ for experiments with wild-type AmnC. For measurements with mutants, a maxiumum concentration of NAD^+^ was 250 μM. Measurements were performed in standard treated capillaries (NanoTemper Technologies) on a Monolith NT.115 system (NanoTemper Technologies) using 50% LED and 80% IRlaser power. The laser on and off times were set to 20 s and 5 s, respectively.

### Preparation of AmnC substrate analogue 2-hydroxymuconate-6-semialdehyde

ACMS was generated enzymatically by catalyzing the di-oxygenation of 3-hydroxyanthranilic acid in an oxygen-saturated buffer with ferrous 3-hydroxyanthranilate 3,4-dioxygenase containing no free transient metal ions, as previously reported. The reaction buffer contained 50 mM MOPS (pH 6.5), Fe(NH4)2SO4, 3-hydroxyanthranilic acid (3-HAA), and 3-hydroxyanthranilate 3,4-dioxygenase (nbaC) protein in a total volume of 2 mL at room temperature. ACMS formation was monitored using an Agilent 8453 diode array spectrophotometer at 360 nm. 2-HMS was generated non-enzymatically by ACMS. ACMS formation was monitored based on the absorbance at 360 nm, then hydrochloric acid was added to adjust the pH to below 3. 2-HMS formation took approximately 25 min, and the absorption maximum was shown to shift irreversibly to 315 nm below pH 3. The solutions were then neutralized to pH 8 with sodium hydroxide after the absorbance at 315 nm stopped increasing, and the absorption maximum shifted to 375 nm.

### Enzyme activity assay

The enzyme activity assay was performed at room temperature on an Agilent 8453 diode array spectrophotometer. After 2-HMS was generated, 1 mM NAD, AmnC-WT, and AmnC mutants were added to catalyze the reaction. Consumption of 2-HMS by AmnC-WT and AmnC mutants was detected by monitoring the decrease in absorbance at 375 nm for 5 min with a 1 min integration time. The absorbance at 375 nm decreased, which is consistent with the results of previous reports.

## Abbreviations

3-HAA: 3-hydroxyanthranilic acid
Pdi: index of polydispersity
ct-FDH: 10-formyltetrahydrofolate dehydrogenase
AmnE: 2-aminomuconate deaminase
2-AMS: 2-aminomuconate-6-semiadehyde
2-AM: 2-aminomuconate
AmnC: 2-aminomuconic 6-=semialdehyde dehydrogenase
AmnBA: 2-aminophenol 1,6-dioxygenase
2-HMS: 2-hydroxymuconate-6-semiadehyde
AmnF: 2-oxopent-4-enoate hydratase
nbaC: 3-hydroxyanthranilate 3,4-dioxygenase
3-HAA: 3-hydroxyanthranilic acid
AmnG: 4-hydroxy-2-oxovalerate aldolase
AmnD: 4-oxalocrotonate decarboxylase
AmnH: acetaldehyde dehydrogenase
ALDH: aldehyde dehydrogenase
AMADHs: aminoaldehyde dehydrogenases
BD: binding domain
CD: catalytic domain
DSF: differential scanning fluorimetry
DLS: dynamic light scattering
Dmax: maximum particle dimension
Tm: melting temparature
NAD^+^: nicotinamide adenine dinucleotide
OD: oligomerization domain
Rg: radius of gyration
RMSD: root-mean-square deviation
SEC: size-exclusion chromatography
SAXS: small-angle X-ray scattering
SSADH: succinic semialdehyde dehydrogenase
ACMS: α-amino; β=carboxymuconate ε-semialdehyde
2-HM: α-hydroxymuconic acid

## Data availability

The Atomic coordinates and structure factors of AmnC was submitted to RCSB Protein Data Bank, with accession code 7BZV. The small angle scattering of X-ray data were submitted to SASBDB small angle scattering biological data bank, with ID 3284 and 3258.

## Acknowledgments

We gratefully acknowledge the core facility staff members of West China Hospital. We thank the staff of the BL19U1 beamline of the National Facility for Protein Science in Shanghai (NFPS) at the Shanghai Synchrotron Radiation Facility for their assistance during data collection.

## Author contributions

D.S., Q.S. designed the study. Q.S. purified and characterized the proteins, cultivated the protein crystals, collected X-ray diffraction data, and managed the biological experiments. Y.C, C.C. solved the crystal structures. Y.C., G.L. collected SAXS data. X.L. assisted with the biological experiments and provided essential data for this study. D.S. and Y.C. took the lead in writing the manuscript. Y.C., Q.S. designed and created all the figures. H.D., Z.Z, and C.Y. contributed to the data analysis and the final version of the manuscript. DS had final approval of the version to be published.

